# Physics and Morphology Constrained Quantitative Susceptibility Based Segmentation of Cerebral Veins

**DOI:** 10.64898/2026.09.24.754280

**Authors:** Yousif C. Al-Khoury, Ali Rezaei, Vanessa Raspa, Nicole Slanina, Hendrale Gresseau, Yiming Xiao, Binu P. Thomas, Claudine J. Gauthier

## Abstract

**Purpose:** Quantitative susceptibility mapping (QSM) provides venous contrast through the paramagnetic susceptibility of deoxyhemoglobin and can be used to estimate oxygen extraction fraction (OEF), a marker of cerebral metabolism. However, cerebral vein segmentation remains challenging due to artifacts, variability across QSM reconstruction methods, and limited sensitivity of conventional vessel-filtering approaches to small cortical veins. This study proposes a physics- and morphology-constrained deep learning framework for cerebral vein segmentation on QSM.

**Methods:** Thirty subjects were manually segmented and used to train an attention-gated U-Net. In addition to supervised Dice and cross-entropy losses, the network incorporated two self-supervised constraints. A physics-informed loss enforced consistency between the measured local field and a field simulated from predicted veins using dipole convolution. A Frangi vesselness loss encouraged anatomically plausible tubular structures. Segmentation performance was evaluated on an independent multi-center cohort and compared with a prior vessel-filtering method and a supervised U-Net baseline. Physiological relevance was assessed by comparing QSM-derived venous OEF with calibrated fMRI-derived OEF.

**Results:** The proposed model achieved superior segmentation performance (centreline Dice = 0.70 ± 0.08) compared with MSVF (0.44 ± 0.09) and the U-Net baseline (0.57 ± 0.14). Improvements were most pronounced in smaller veins, resulting in greater vessel continuity. Venous OEF derived from the proposed framework demonstrated stronger agreement with calibrated fMRI-derived OEF than the previous approach.

**Conclusion:** Combining supervised learning with physics- and morphology-based constraints improves vein segmentation and enhances the sensitivity of QSM-derived OEF measurements. Code and model weights are publicly available at https://github.com/QuantitativePhysiologyImagingLab/VeinSeg.

## 1 Introduction

The venous network is essential to bring deoxygenated blood back to the lungs after it has circulated through the organs. Venous alterations have been reported in numerous brain pathologies, including cerebrovascular disease, dementia, multiple sclerosis, diabetes, and aging [1, 2]. These conditions impact venous morphology and hemodynamics, making in vivo quantification of venous structure and function valuable for assessing cerebral vascular and metabolic physiology [3, 4]. The cerebral vein network provides important physiological information about tissue metabolism, as venous blood reflects local oxygen extraction and consumption [5]. However, imaging cerebral veins remains challenging. MRI approaches such as time-of-flight and MR angiography primarily target arterial structures and have limited sensitivity to small cortical veins [6]. They also require long scan times and are prone to motion artifacts, while contrast-enhanced variants additionally require intravenous gadolinium [7, 8]. Furthermore, such datasets are far less common in public neuroimaging and clinical databases compared to gradient-echo (GRE) acquisitions, which are routinely collected across clinical and research settings. Quantitative susceptibility mapping (QSM) is a post-processing technique applied to GRE data that estimates the bulk magnetic susceptibility of tissue [9, 10]. QSM is sensitive to iron deposition and demyelination, making it relevant to neurodegeneration, and additionally provides strong contrast for cerebral veins due to the paramagnetic susceptibility of deoxyhemoglobin [11–15].

One major application of venous QSM is oxygen extraction fraction (OEF) estimation [14, 16, 17], a metabolic marker linked to both cerebral perfusion and oxidative metabolism, reflecting the balance between oxygen delivery and consumption [18–20]. OEF can be estimated from the susceptibility difference between paramagnetic deoxyhemoglobin in venous blood and surrounding tissue, enabling non-invasive assessment of cerebral oxygen utilization [21, 22].

OEF is tightly regulated under normal conditions but is altered in cerebrovascular and neurodegenerative diseases, making it a valuable biomarker of impaired oxygen supply and metabolic dysfunction [18]. This approach exploits the magnetic susceptibility difference between venous blood, where deoxygenated hemoglobin is paramagnetic, and surrounding tissue, which is weakly diamagnetic [21]. The resulting susceptibility contrast is directly related to venous oxygen saturation and hematocrit, both of which determine the concentration of deoxyhemoglobin [22]. While established techniques such as oxygen-15 positron emission tomography (^15^O-PET), calibrated functional MRI (fMRI), quantitative BOLD (qBOLD), and T_2_-Relaxation-Under-Spin-Tagging (TRUST) MRI can also assess OEF, they require radioactive tracers, respiratory manipulations, specialized acquisitions, or provide only limited spatial specificity [23–27]. In contrast, venous QSM provides a regionally resolved, gas-free alternative that can be derived from routinely acquired GRE data [28, 29].

Several prior studies have demonstrated meaningful correspondence between venous QSM-derived OEF and established reference measures. Kudo et al. [30] demonstrated agreement between MRI-based venous oxygenation measurements and ^15^O-PET OEF, supporting the physiological validity of venous QSM relative to the current reference standard for cerebral OEF imaging. Similarly, Fan et al. [19] reported significant correlations between susceptibility-based venous OEF and calibrated fMRI-derived OEF in healthy volunteers. Collectively, these findings suggest that venous QSM has the potential to provide physiologically meaningful measurements of cerebral oxygen metabolism using standard and widely available MRI acquisitions.

However, current QSM-based venous segmentation methods remain limited in their ability to comprehensively capture all cerebral veins, especially those with smaller diameters. Accurate characterization of these finer vessels is particularly important for OEF estimation, as these capture more local metabolism than large draining veins such as the sagittal sinus [31]. Existing approaches to segment veins on QSM images include threshold-based methods [30], but these struggle to pick up the varying contrasts seen in veins. Other approaches are based on Hessian or Frangi vesselness filters [32, 33], which rely on hand-tuned parameters and often degrade in low-contrast or noisy regions, leading to inconsistent detection of smaller structures. More recently, Kim et al. [34] developed a susceptibility-based vessel segmentation by combining vessel filtering with seed generation and region growing, utilizing 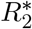 and *χ*-separation information. Despite improved performance, the method remains dependent on handcrafted vesselness filters, region-growing heuristics, and subject-specific parameter tuning, and *χ*-separation tailored acquisitions.

Deep learning has transformed vessel segmentation in other imaging modalities, including angiography and retinal imaging, but remains largely unexplored for venous segmentation on QSM. Existing vessel foundation models such as VesselFM [35] and VesselBoost [36] have primarily been trained on angiographic datasets that emphasize arterial anatomy and may not generalize to the unique contrast mechanisms of QSM. To our knowledge, no prior method has combined deep learning with physics-based constraints for QSM venous segmentation. We therefore propose a physics- and morphology-constrained deep learning framework that leverages susceptibility-based contrast mechanisms and vascular priors, improving generalization beyond the training data while improving sensitivity to small vessels.

## 2 Methods

### 2.1 Study Cohorts

Five datasets were used for segmentation model development, segmentation evaluation, and down-stream physiological validation (Table 1). The training cohort consisted of healthy controls (Dataset 1), individuals with coronary artery disease, type-2 diabetes, and controls (Dataset 3), and an AD subtyping cohort (Dataset 5), acquired across 7T and 3T field strengths to increase variability in venous contrast, susceptibility appearance, and physiological state. Segmentation performance was evaluated on an independent multi-center cohort comprising healthy controls (Dataset 1), an independent coronary artery disease (CAD) subject (Dataset 3), a cohort including Parkinson’s disease, intracranial calcifications, and controls (Dataset 6), and calculation of susceptibility through multiple orientation sampling (COSMOS) data (Dataset 4).

**Table 1:**
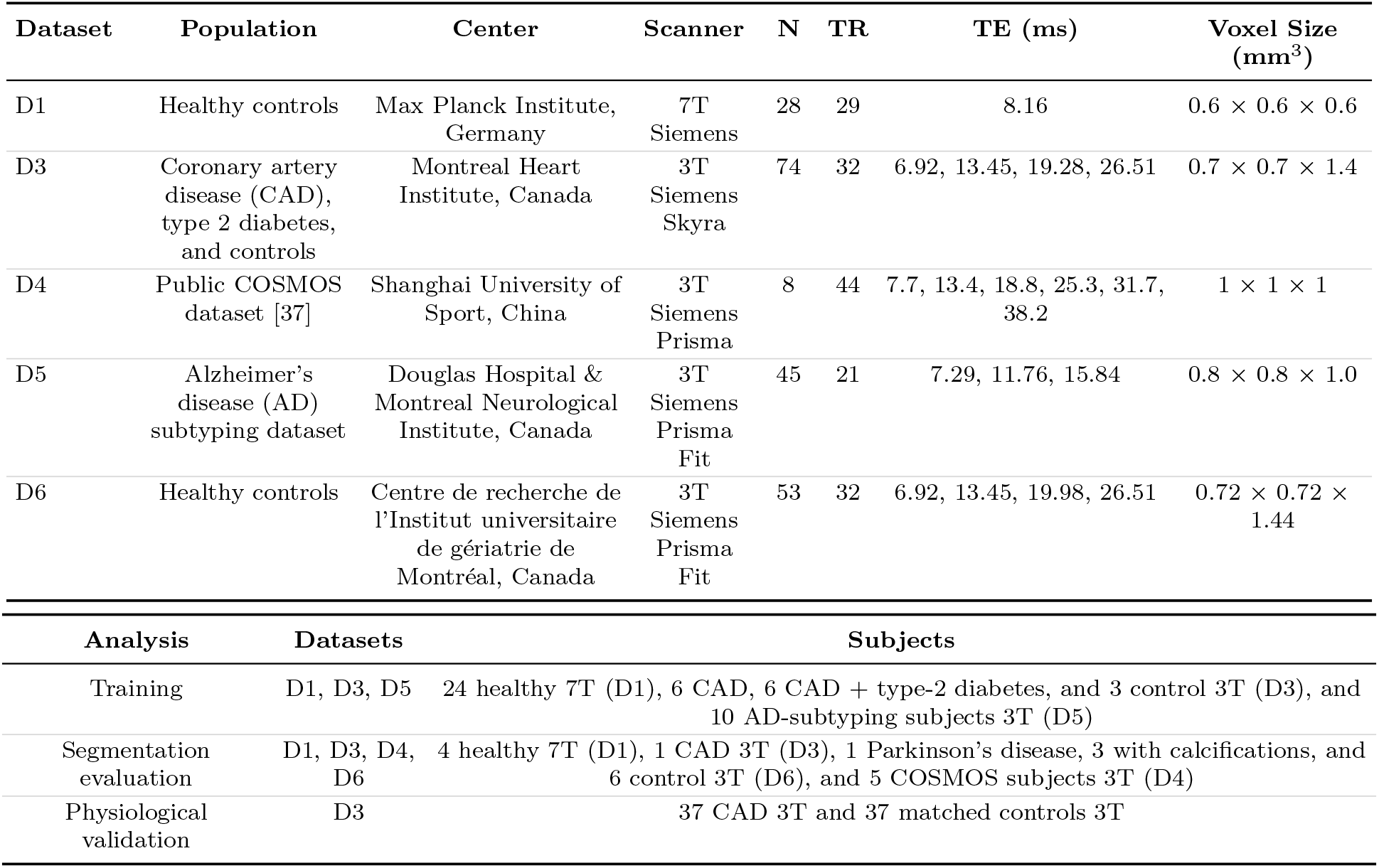
Summary of study cohorts, acquisition parameters, and dataset allocation.

The CAD cohort (Dataset 3) was additionally used for downstream physiological validation of venous OEF measurements. In addition to multi-echo GRE imaging, this cohort underwent calibrated fMRI acquisitions to enable estimation of OEF using the gas calibration model (GCM). Flow compensation was applied on the first GRE echo to minimize phase artifacts arising from slow venous flow. Venous OEF derived from the proposed segmentation framework was subsequently compared against GCM-derived OEF measurements acquired during the same imaging session.

Five subjects from a publicly available COSMOS dataset (Dataset 4) were included for anatomical validation. Because COSMOS combines multiple head orientations to obtain highly accurate susceptibility estimates, it provided a strong anatomical reference to evaluate venous structures [37, 38]. All studies were approved by their local research ethics committees in accordance with the Declaration of Helsinki.

### 2.2 QSM Processing

For all acquisitions, phase images were unwrapped using rapid open-source minimum spanning tree algorithm (ROMEO) [39], and background field removal was performed using regularization enabled sophisticated harmonic artifact reduction for phase data (RESHARP) [40]. QSM images were subsequently generated using multiple reconstruction approaches for each acquisition, including total variation regularized QSM (TGV-QSM) [41], morphology enabled dipole inversion (MEDI) [42], iterative least-squares (iLSQR) [43], STAR-QSM [44], and L1-regularized reconstructions [45]. TGV-QSM was selected for manual annotation due to its superior venous contrast and vessel delineation relative to the alternative methods. All subsequent venous OEF measurements were computed using MEDI reconstructions due to their stable venous susceptibility values [46].

Venous structures were manually segmented on TGV-QSM images in all three orthogonal planes by trained annotators (N.S., H.G., S.C., T.K., H.M.) under the supervision of (A.R.), who has extensive experience identifying cerebral veins on QSM. An initial subset of 15 subjects was manually segmented entirely from scratch and used to train an early version of the segmentation model. The resulting predictions from this preliminary model were then manually refined across all slices and planes to generate annotations for the remaining 15 subjects, including six 3T scans. The 30 binary segmentations served as ground-truth labels for supervised training and evaluation. These same manual segmentations were subsequently transferred to the alternative QSM reconstructions and refined by retaining only voxels with positive susceptibility values on the corresponding reconstruction, accounting for reconstruction-specific differences in venous contrast and artifact behavior [46]. Including all QSM reconstruction variants, a total of 150 images were used during training.

### 2.3 Calibrated fMRI Processing

Calibrated fMRI was acquired as part of Dataset 3 in all participants. GCM-based OEF was estimated from dual-echo pseudo-continuous arterial spin labeling (pCASL) data acquired during controlled respiratory manipulations, targeting sequential hypercapnic (+5 mmHg above baseline end-tidal CO_2_), isocapnic hyperoxic (+150 mmHg above baseline end-tidal O_2_), and combined hyperoxia–hypercapnia conditions, each sustained for 2 min and separated by 2 min of room air inhalation, using the RespirAct− system (Thornhill Research, Toronto, Canada). End-tidal partial pressures of CO_2_ and O_2_ were used as proxies for arterial gas concentrations [47]. Perfusion-weighted time series were derived from the first echo via surround subtraction, and BOLD-weighted signals from the second echo via surround addition. OEF was subsequently estimated from the BOLD and ASL responses to the respiratory challenges using the GCM [25].

### 2.4 Network Architecture

This model was trained on a customized nnU-Net framework [48], utilizing its preprocessing pipeline but replacing the default 3D U-Net with a customized attention-gated U-Net backbone [49]. The architecture inserts attention gates on the skip connections and a prior-gating block on the input channels.

The first channel corresponds to the anatomical QSM image, while the remaining channels represent priors to guide training onto specific regions, specifically the local field after background removal, and a Frangi vesselness map computed by filtering the QSM. In addition to implementing the attention gates described in Oktay et al. [49], these prior channels are processed by a prior gate module, which learns a spatial modulation field indicating how strongly the network should attend to each voxel based on the confidence provided by the priors. The modulation is applied to the anatomical image, which amplifies the features extracted from the QSM image in locations where the priors suggest higher importance to improving the model performance (vessel regions), while leaving other regions unchanged. The modulated image is then concatenated with the untouched prior channels and passed into the attention-gated U-Net. This mechanism encourages early feature extraction to focus on venous structures without rigidly imposing the priors on the output.

Additionally, to improve robustness across different QSM reconstruction methods and field strengths, the network was conditioned using feature-wise linear modulation (FiLM) [50]. Learnable embeddings were assigned to each QSM reconstruction method (TGV, MEDI, iLSQR, L1, and STAR-QSM) and field strength (3T or 7T). These embeddings were used to modulate intermediate feature maps throughout the encoder and bottleneck layers via channel-wise scaling and shifting operations. This allowed the network to adapt its internal feature representations according to differences in image contrast, noise characteristics, and susceptibility appearance between reconstruction pipelines and field strengths, while still sharing a common set of network weights across all inputs. The model architecture is visualized in Figure 1.

**Figure 1:**
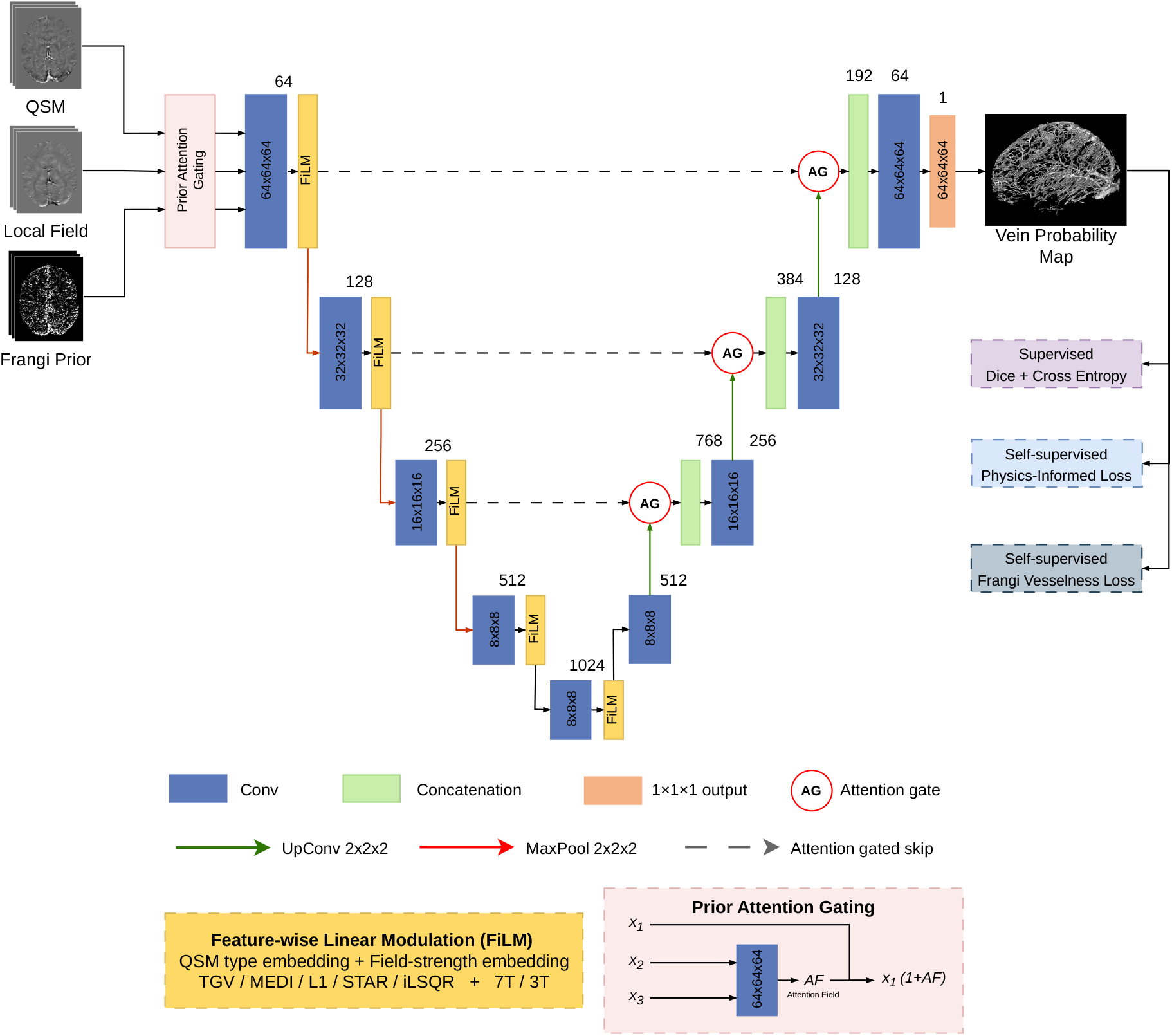
Overview of the proposed physics-guided segmentation architecture. The network receives three input channels: the QSM image (channel 1), the measured local field after background removal (channel 2), and a Frangi vesselness map computed from the QSM (channel 3). A prior-gate module learns a spatial modulation field from channels 2 and 3, which amplifies anatomical QSM features in vessel-likely regions before concatenation with the prior channels. The resulting input is processed by an attention-gated U-Net conditioned via FiLM embeddings encoding reconstruction method and field strength. During training, the predicted vein probability map drives three loss terms: the standard overlap-based supervised segmentation loss, a physics-informed local field loss that enforces consistency between the dipole-simulated field from predicted venous susceptibility and the measured local field, and a Frangi-based vesselness loss that encourages tubular vascular morphology consistent with the QSM-derived prior.

### 2.5 Training Strategy

The model was trained in three sequential stages. Each stage was run for 1000 epochs, at which point the next loss term was introduced.

#### Stage 1: Baseline Cross-entropy and Dice

Training began with standard cross-entropy and soft Dice loss on manual vein labels, reflecting the commonly adopted baseline in deep-learning based segmentation literature. This stage was performed twice, first using a vanilla U-Net (baseline architecture provided by nnUNet [48]), serving as a reference baseline, and then using the proposed attention- and prior-gated U-Net. All subsequent stages were trained using the attention-based architecture.

#### Stage 2: Physics-constrained training

A dipole convolution forward model was used to enforce physical consistency between the predicted venous susceptibility and the local field *B*_local_ (provided as the second input channel).

#### Stage 3: Frangi shape prior

A Hessian-based vesselness (Frangi) loss term was added to Stage 2, encouraging tubular and anatomically plausible vein structures that align with the tubular morphology visible in the QSM image (third input channel).

### 2.6 Physics-Informed Local Field Loss

Let *χ*_tissue_(**r**) denote the reconstructed tissue susceptibility in ppm (channel 1), and *B*_meas_(**r**) the measured local magnetic field after background field removal using RESHARP (channel 2) [40]. The effective venous blood susceptibility *χ*_blood_ (ppm) was estimated for each reconstruction technique and field strength by averaging susceptibility values within the corresponding group’s ground-truth venous segmentation. From the network output, a binary vein mask is derived:

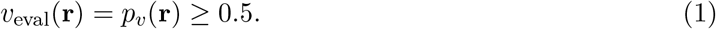

This vein mask is then used to construct a susceptibility distribution corresponding to deoxygenated venous blood:

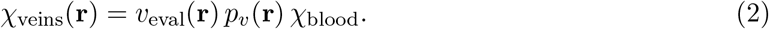

This formulation assumes that venous susceptibility sources are a dominant contributor to the local magnetic field perturbations observed in QSM phase data, primarily due to the strong paramagnetic susceptibility of deoxyhemoglobin. Consequently, the local field generated from the predicted venous susceptibility distribution should approximate the measured local field obtained following RESHARP background field removal.

The predicted vein local field *B*_pred_ is computed using a standard Fourier-domain dipole convolution. Let **k** = (*k*_*x*_, *k*_*y*_, *k*_*z*_) denote spatial frequencies (constructed from the voxel sizes *s*_*x*_, *s*_*y*_, *s*_*z*_), and 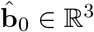 the unit vector along the main magnetic field direction *B*_0_. The dipole kernel is:

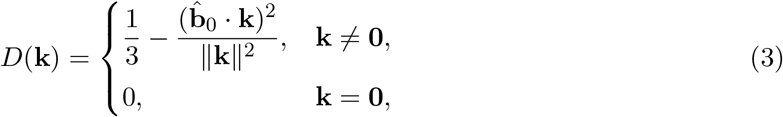

and the predicted field is:

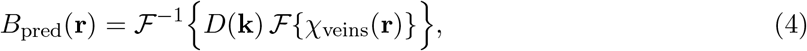

where *ℱ* denotes the 3D fast Fourier Transform (FFT). A simplified pipeline of the physics-constrained loss is visualized in Figure 2.

**Figure 2:**
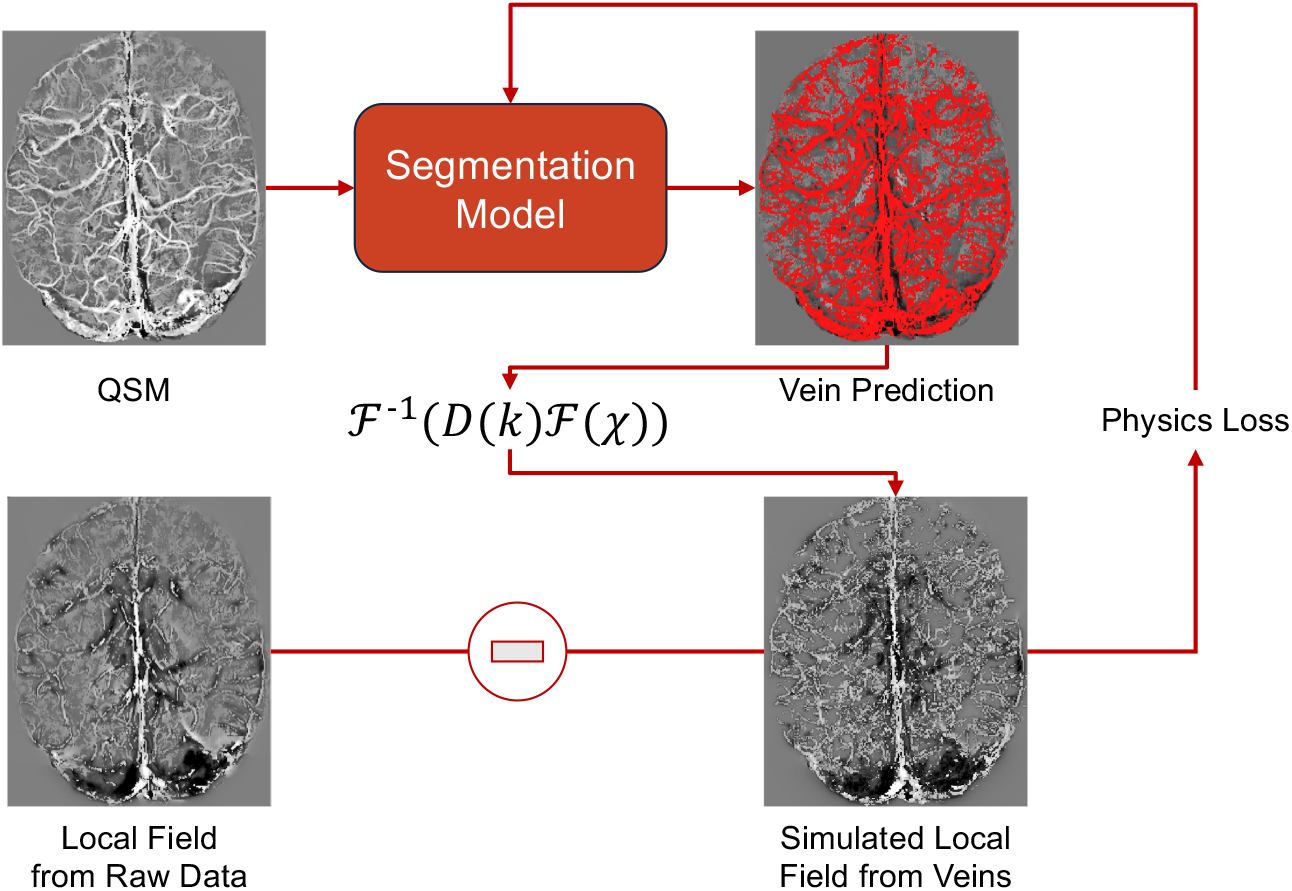
Physics-informed training framework. The segmentation network predicts a venous mask from QSM. A dipole forward model is then applied to the predicted veins to generate a simulated local field, which is compared with the measured local field obtained from the raw phase data. The resulting physics loss constraints the model to enforce consistency between the predicted venous anatomy and the susceptibility-induced magnetic field perturbations.

#### 2.6.1 Whole-Brain Weighted Mean Absolute Error

Magnetic field perturbations from veins are not always local to the veins due to the strong paramagnetic nature of deoxyhemoglobin, causing the local field values in nearby tissue to be affected by veins. Therefore, a loss was constructed to consider the whole brain when comparing local fields. However, larger perturbations need to be weighted more as the larger values reflect proximity to vein regions which are more relevant to guiding the model. The loss computes a weight map where regions with larger measured local field perturbations contribute more strongly to the optimization objective. The weight map is clamped to a maximum value of 5 to prevent a small number of high-field voxels from dominating the loss. This cap preserves increased weighting near venous susceptibility sources while maintaining contributions from the surrounding tissue field pattern and improving training stability.

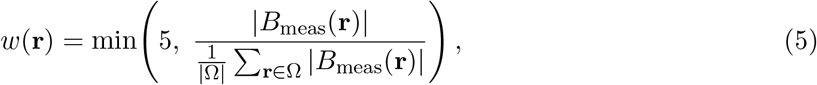

and the weighted mean absolute error is defined over the brain:

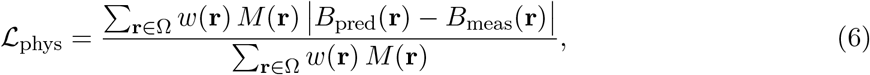

where Ω represents voxels inside the brain.

#### 2.6.2 Vein-Restricted MAE and Sign Hinge Loss

Over the vein evaluation region Ω_vein_, define:

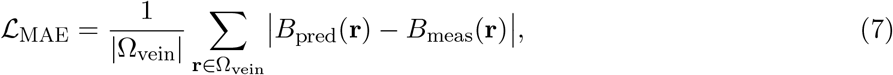

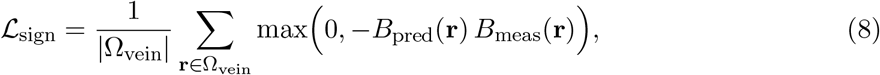

where Ω_vein_ represents voxels within the predicted vein. These losses penalize large magnitude discrepancies and wrong field polarity, respectively.

#### 2.6.3 Combined Physics Loss

The physics-informed loss is a weighted sum of these components:

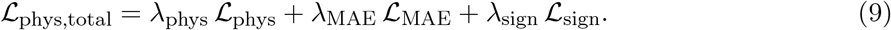

In Stage 2, the total loss becomes:

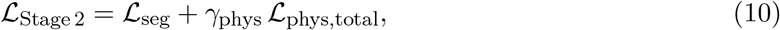

where *γ*_phys_ controls the overall strength of the physics constraint.

### 2.7 Frangi-Based Vessel Shape Loss

To encourage anatomically plausible vessel morphology, a Frangi vesselness filter [33] was used to derive tubular structure priors from both the QSM image and the predicted vein segmentation. Briefly, the Frangi filter computes a multiscale vesselness response from the eigenvalues of the Hessian matrix, producing high responses in elongated tubular structures and low responses in blob-like or planar regions.

The multiscale vesselness prior derived from the QSM image was defined as:

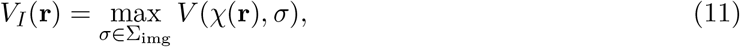

where *V* (·, *σ*) denotes the Frangi vesselness response at scale *σ*. The resulting vesselness map *V*_*I*_ was provided as an input channel and additionally served as an anatomical prior during training.

Similarly, vesselness was computed from the hard-thresholded prediction:

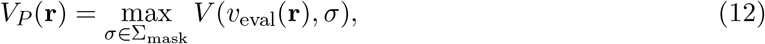

and subsequently rescaled such that its peak response within the gated region matched that of *V*_*I*_.

To focus the loss onto vessel-like regions, a gate was constructed from the image vesselness:

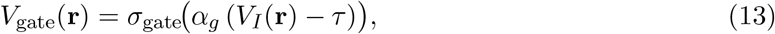

where *σ*_gate_ denotes the sigmoid function, *α*_*g*_ controls its steepness, and *τ* is a small threshold. *V*_gate_ was subsequently dilated using a 3×3×3 max-pooling operation to bridge small discontinuities. The active region Ω consisted of brain voxels where *V*_gate_ *>* 0.51.

#### 2.7.1 Vesselness Term

To encourage the network to assign high probability to regions that are both tubular and consistent with the image prior:

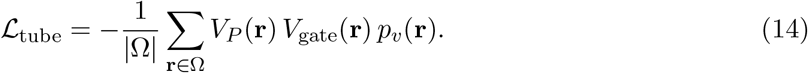

#### 2.7.2 Completion Term

To penalize predictions that were less tubular than suggested by the image prior, a one-sided hinge loss was introduced. Because predictions are binary masks rather than continuous images, they can naturally exhibit higher vesselness responses than the QSM-derived prior. The hinge therefore activates only when the prediction vesselness falls below the image vesselness:

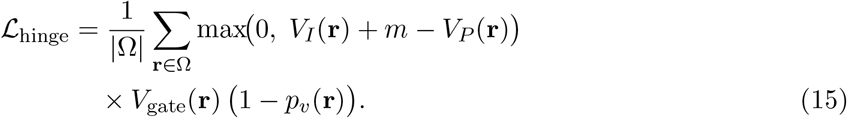

with margin *m* = 0.1. Minimizing this term encourages *p*_*v*_ to increase in regions where the image Frangi prior indicates vessel-like structure but the prediction remains incomplete.

#### 2.7.3 Combined Frangi Loss

The two Frangi terms serve complementary roles. The vesselness term globally encourages tubular and vessel-like predictions within regions supported by the image prior, promoting continuity and coherent vascular morphology. In contrast, the hinge term acts specifically on regions where the QSM-derived vesselness exceeds the prediction vesselness, thereby encouraging completion of missing or fragmented vessel segments without penalizing predictions that are already more tubular than the image prior:

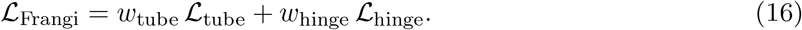

In Stage 3, the total loss being minimized is:

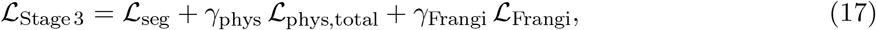

where *γ*_Frangi_ controls the strength of the Frangi loss.

### 2.8 Performance Evaluation

Segmentation performance was evaluated globally across the whole brain and separately within each cortical lobe. The caudate, putamen and thalamus were excluded due to high iron content, which reduces the reliability of venous annotation, while the cerebellum was excluded because of susceptibility artifacts associated with nearby air–tissue interfaces. To evaluate the contribution of each proposed component, an ablation analysis was performed comparing the baseline vanilla U-Net, the physics-constrained model, the Frangi-constrained model, and the full combined framework.

Global segmentation performance was quantified using the Dice similarity coefficient, precision, recall, and average symmetric surface distance (ASSD) [51]. Because global Dice similarity is dominated by larger venous structures and may therefore obscure improvements in smaller cortical veins, an additional diameter-stratified analysis was performed to specifically assess sensitivity to fine venous anatomy. Vein segmentations were first skeletonized to obtain vessel centrelines, after which a Euclidean distance transform was computed to estimate the local vessel radius from the distance between each centreline voxel and the vessel boundary. Vessel diameter was subsequently defined as twice the local radius. Ground-truth veins were then grouped into diameter categories of *<* 1 mm, *<* 2 mm, and *>* 4 mm, and Dice scores were computed separately within each category. This analysis was intended to determine whether the proposed physics and Frangi constraints preferentially improved recovery of smaller venous structures, which are more susceptible to partial-volume effects and are typically under-segmented by conventional approaches.

### 2.9 Venous OEF

OEF was estimated from venous susceptibility using a biophysical relationship between magnetic susceptibility and venous oxygen saturation [12, 14, 31, 32]. For each venous QSM voxel, OEF was computed as:

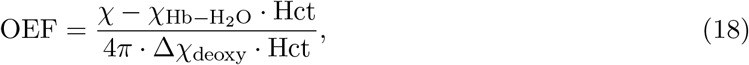

where *χ* is the calibrated QSM value in ppm, Hct = 0.40 is the assumed hematocrit, Δ*χ*_deoxy_ = 0.18 ppm represents the susceptibility contribution of fully deoxygenated hemoglobin, and 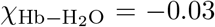 ppm accounts for the hemoglobin–water susceptibility offset [12, 14]. This formulation assumes near-complete arterial oxygen saturation under normal physiological conditions [14].

To reduce partial-volume contamination, susceptibility values were corrected using the venous probability map produced by the proposed segmentation model. OEF values derived using the proposed framework were compared against OEF estimated using the multiscale vessel filtering (MSVF) approach, which identifies venous structures using a recursive ridge filtering method applied on the QSM image [32]. Both approaches were evaluated against GCM-derived OEF measurements, which served as the reference standard for non-invasive cerebral oxygenation assessment.

Region-wise OEF measurements were obtained using anatomical masks derived from the MNI structural atlas registered to the QSM space [52, 53]. Subcortical and susceptibility artifact-prone regions, including the caudate, cerebellum, putamen, and thalamus, were excluded from whole-brain cortical OEF estimation due to the effects of high susceptibility regions, such as iron-rich deep gray matter, on venous susceptibility which would contaminate OEF measurements.

## 3 Results

### 3.1 Segmentation Performance

Vein segmentation performance on TGV-QSM is summarized in Table 2 and visualized in Figure 3. The proposed full model achieves the best overall performance across the evaluated methods. Compared with the vanilla U-Net, the full model improves Dice, clDice, recall, and surface distance error. The MSVF baseline yields lower performance than the learning-based approaches across all metrics. Foundation model approaches, namely VesselFM, were also evaluated but excluded from the comparison because the foundation model was not trained on QSM data and failed to reliably recover any venous structures [35].

**Table 2:** Segmentation performance on TGV-QSM. Quantitative comparison between MSVF, Vanilla U-Net, and the proposed model across *n* = 21 held-out cases.

| Method | DSC | clDSC | Precision | Recall | ASSD (mm) |
| --- | --- | --- | --- | --- | --- |
| MSVF | $0.43 \pm 0.09$ | $0.44 \pm 0.09$ | $0.37 \pm 0.09$ | $0.53 \pm 0.10$ | $1.59 \pm 0.49$ |
| Vanilla U-Net | $0.61 \pm 0.12^{**}$ | $0.57 \pm 0.14^{**}$ | $0.56 \pm 0.10^{**}$ | $0.69 \pm 0.15^{**}$ | $1.94 \pm 3.81^{**}$ |
| **Proposed** | **$0.74 \pm 0.06^{**}$** | **$0.70 \pm 0.08^{**}$** | **$0.70 \pm 0.11^{**}$** | **$0.80 \pm 0.05^{**}$** | **$0.72 \pm 0.33^{**}$** |

\* $p < 0.05$ versus MSVF (Wilcoxon signed-rank test, $n = 21$ ).
\*\* $p < 0.01$ versus MSVF (Wilcoxon signed-rank test, $n = 21$ ).

**Figure 3:**
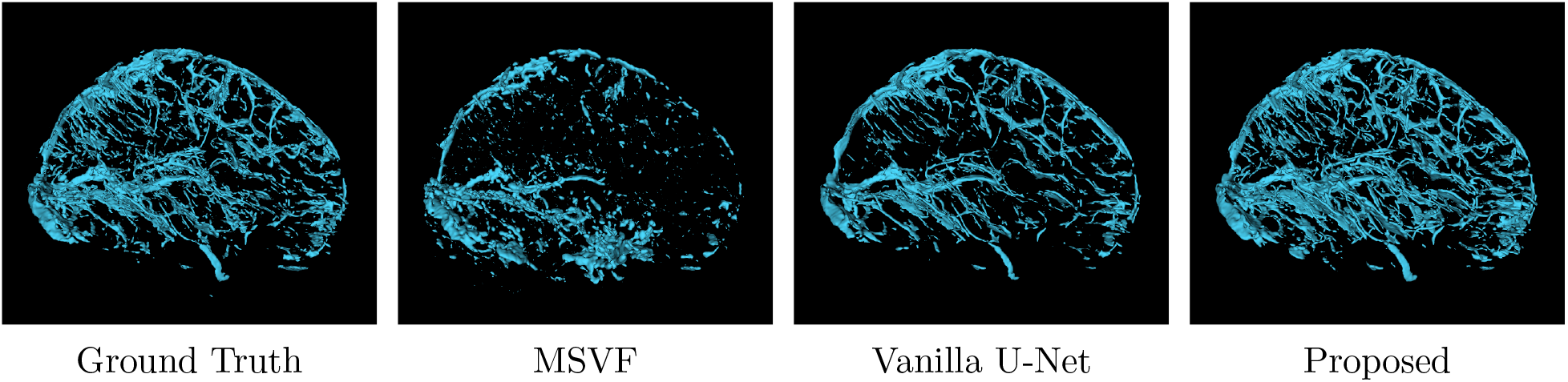
Example 3D visualization of cerebral vein segmentation on a 7T QSM image. From left to right: manual ground truth, MSVF, Vanilla U-Net, and the proposed model. The proposed model demonstrates improved recovery and continuity of smaller venous structures compared with the baseline approaches.

### 3.2 Ablation Study

The ablation study is summarized in Table 3A. Replacing the vanilla U-Net with the attention-based backbone does not substantially change performance. Adding the physics loss increases recall and reduces ASSD. Adding the Frangi loss further increases recall, while the full model achieves the best performance across all five metrics, including the highest Dice and the lowest ASSD. Diameter-stratified Dice is shown in Table 3B. The *<* 1 mm category includes only 7T subjects, as sub-millimeter veins are not visible at 3T resolution due to the lower resolution in the slice direction. The full model achieved the highest Dice for both vessels below 1 mm and vessels below 2 mm, indicating that the combined constraints provide the greatest benefit for the smallest, most difficult-to-segment veins. All models showed more comparable performance for vessels larger than 4 mm, although the vanilla U-Net remained modestly lower than the three constrained configurations.

**Table 3:** Quantitative evaluation of the proposed framework. (A) Ablation study evaluating the contribution of the Physics and Frangi losses (*n* = 21 held-out cases). (B) Diameter-stratified Dice analysis demonstrating performance across vessel calibers (*n* = 13 for *<* 2 mm and *>* 4 mm; *n* = 4 for *<* 1 mm, 7T only). Values are mean ± SD.

| **(A) Ablation Study** |  |  |  |  |  |
| --- | --- | --- | --- | --- | --- |
| Model | DSC | clDSC | Precision | Recall | ASSD (mm) |
| Vanilla U-Net | $0.61 \pm 0.12$ | $0.57 \pm 0.14$ | $0.56 \pm 0.10$ | $0.69 \pm 0.15$ | $1.94 \pm 3.81$ |
| Physics loss | $0.70 \pm 0.05^{**}$ | $0.64 \pm 0.09^{**}$ | $0.68 \pm 0.07^{**}$ | $0.72 \pm 0.06$ | $0.89 \pm 0.39^{**}$ |
| Frangi loss | $0.70 \pm 0.06^{**}$ | $0.66 \pm 0.08^{**}$ | $0.64 \pm 0.08^{**}$ | $0.78 \pm 0.05^{**}$ | $0.83 \pm 0.39^{**}$ |
| **Proposed** | **$0.74 \pm 0.06^{**}$** | **$0.70 \pm 0.08^{**}$** | **$0.70 \pm 0.11^{**}$** | **$0.80 \pm 0.05^{**}$** | **$0.72 \pm 0.33^{**}$** |
| **(B) Diameter-Stratified Dice** |  |  |  |  |  |
| Model | $< 1$ mm<sup>†</sup> | $< 2$ mm | $> 4$ mm | | |
| Vanilla U-Net | $0.22 \pm 0.03$ | $0.38 \pm 0.20$ | $0.83 \pm 0.07$ | | |
| Physics Only | $0.56 \pm 0.01$ | $0.39 \pm 0.18$ | **$0.91 \pm 0.03^{**}$** | | |
| Frangi Only | $0.60 \pm 0.02$ | $0.45 \pm 0.17^{**}$ | $0.88 \pm 0.05^{**}$ | | |
| **Proposed** | **$0.62 \pm 0.02$** | **$0.50 \pm 0.15^{**}$** | $0.90 \pm 0.04^{**}$ | | |

\* $p < 0.05$ versus Vanilla U-Net baseline (Wilcoxon signed-rank test).
\*\* $p < 0.01$ versus Vanilla U-Net baseline (Wilcoxon signed-rank test).
<sup>†</sup> Results for vessels $< 1$ mm computed using only the 7T cohort ( $n = 4$ ). Statistical significance testing was not performed for this category due to insufficient sample size.

### 3.3 Generalization Across QSM Reconstructions

Table 4 reports model performance on the test subjects when the QSM input was replaced with four alternative reconstruction algorithms. Performance remained relatively consistent across three of the four reconstruction methods, with Dice scores ranging from 0.66 ± 0.08 (iLSQR) to 0.73 ± 0.06 (MEDI and STAR-QSM), while iLSQR trailed the other methods across most metrics. clDice followed a similar pattern, ranging from 0.62 (iLSQR) to 0.68 (MEDI). MEDI achieved the highest precision (0.67 ± 0.09), while recall was essentially uniform across all four reconstruction methods (0.81–0.82), indicating that reconstruction method primarily influenced the specificity rather than the sensitivity of the predicted venous maps. ASSD varied more substantially between methods: STAR-QSM achieved the lowest surface distance error (0.63 ± 0.32 mm), whereas iLSQR showed the highest (0.75 ± 0.38 mm), indicating greater spatial offset of the predicted venous structures despite comparable overlap-based performance to MEDI and L1.

**Table 4:** Full model performance across QSM reconstruction methods (*n* = 16 for MEDI; *n* = 14 for iLSQR, L1, and STAR-QSM).

| QSM | DSC | clDSC | Precision | Recall | ASSD (mm) |
| --- | --- | --- | --- | --- | --- |
| MEDI | $0.73 \pm 0.06$ | $0.68 \pm 0.05$ | $0.67 \pm 0.09$ | $0.82 \pm 0.02$ | $0.71 \pm 0.33$ |
| iLSQR | $0.66 \pm 0.08$ | $0.62 \pm 0.09$ | $0.56 \pm 0.10$ | $0.82 \pm 0.02$ | $0.75 \pm 0.38$ |
| L1 | $0.70 \pm 0.06$ | $0.66 \pm 0.08$ | $0.61 \pm 0.08$ | $0.82 \pm 0.02$ | $0.68 \pm 0.34$ |
| STAR-QSM | $0.73 \pm 0.05$ | $0.67 \pm 0.06$ | $0.66 \pm 0.06$ | $0.81 \pm 0.03$ | $0.63 \pm 0.32$ |

### 3.4 Venous QSM-Based OEF

The proposed method yielded significant positive correlations with GCM-derived OEF in the parietal lobe (*r* = +0.42, *p <* 0.001) and occipital lobe (*r* = +0.40, *p <* 0.001), with a significant association also observed for whole-brain OEF (*r* = +0.34, *p <* 0.01). The frontal (*r* = +0.12) and temporal (*r* = +0.19) lobes did not reach significance. In contrast, the MSVF approach showed only weak and non-significant correlations across all regions, including the whole brain (*r* = +0.07), frontal lobe (*r* = −0.04), parietal lobe (*r* = +0.10), temporal lobe (*r* = +0.06), and occipital lobe (*r* = +0.09) (Figure 4).

**Figure 4:**
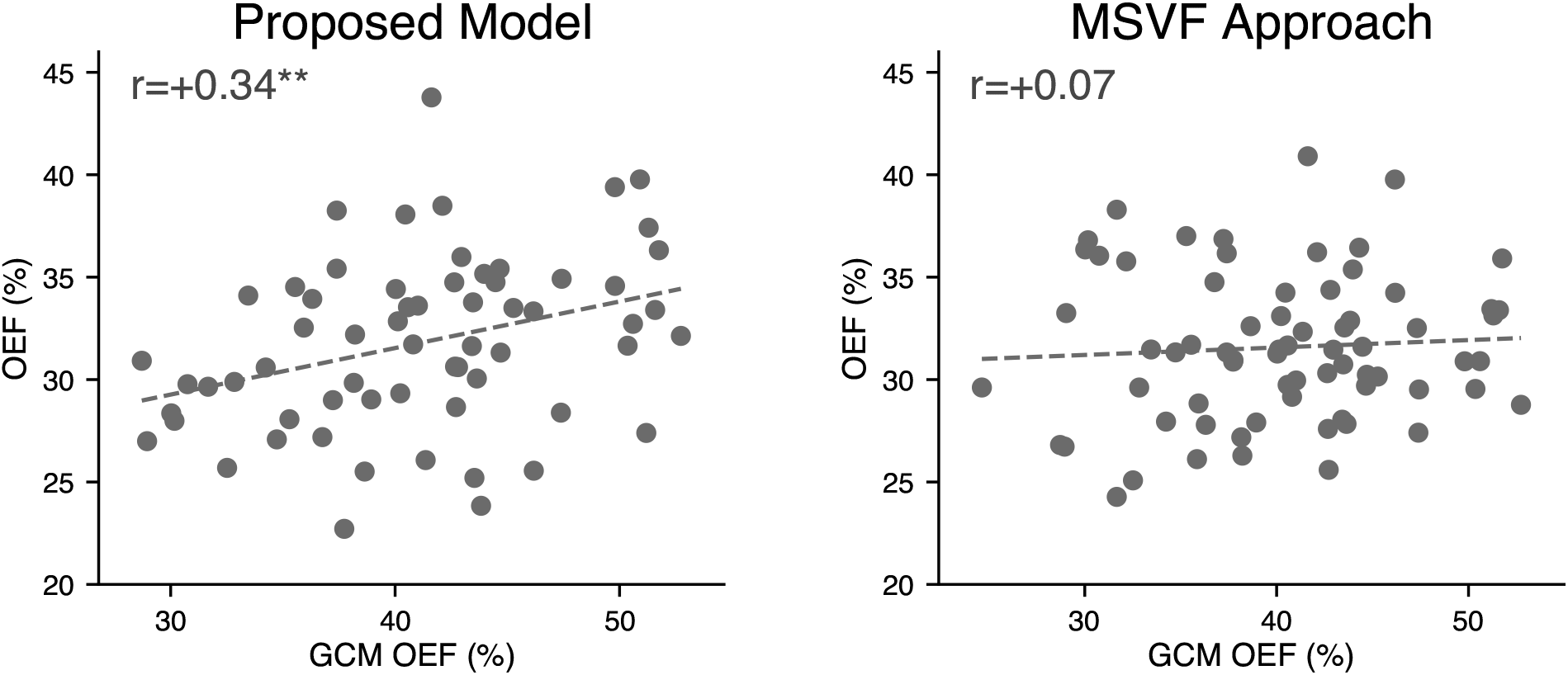
Whole-brain correlation between venous QSM-based OEF and GCM-derived OEF for the proposed model (left) and the MSVF approach (right). Each point represents one subject (*n* = 74). The proposed model demonstrated a stronger and statistically significant association with GCM-derived OEF (*r* = 0.34, *p <* 0.01) compared to the MSVF approach (*r* = 0.07, *p >* 0.1). Improved recovery of small cortical veins provides more physiologically representative venous sampling, enhancing the sensitivity of QSM-derived OEF measurements to underlying cerebral oxygen metabolism.

## 4 Discussion

This work introduced a physics-constrained deep learning framework for cerebral vein segmentation on QSM. Compared to conventional approaches, the proposed method demonstrated improved recovery of venous structures, particularly within smaller cortical veins that are typically challenging to delineate using threshold- or vesselness-based methods alone. Importantly, the benefits of the proposed framework extended beyond voxelwise segmentation metrics and influenced downstream physiological analyses. Venous OEF measurements demonstrated stronger correspondence with OEF values derived from a calibrated fMRI approach, suggesting that improved recovery of smaller cortical veins enhances the physiological sensitivity of venous OEF. Given that venous OEF can be estimated from routinely acquired GRE data, these findings may improve accessibility of cerebral oxygen metabolism quantification.

The ablation study demonstrated that the addition of physics- and Frangi-based constraints improved sensitivity and network completeness compared to the baseline U-Net model, with the largest effects observed in smaller vessels. The physics loss improved spatial agreement between predicted and reference venous boundaries, reflected by reduced surface distance error, while also increasing sensitivity to venous structures. The Frangi loss further enhanced sensitivity by encouraging tubular predictions, although at the cost of reduced precision (Figure 5), indicating that vesselness optimization alone increases false-positive predictions. This highlights the complementary roles of the two constraints: the Frangi term promotes detection of vessel-like structures, while the physics constraint suppresses anatomically implausible predictions and limits oversegmentation. The full model provided the strongest balance between these effects.

**Figure 5:**
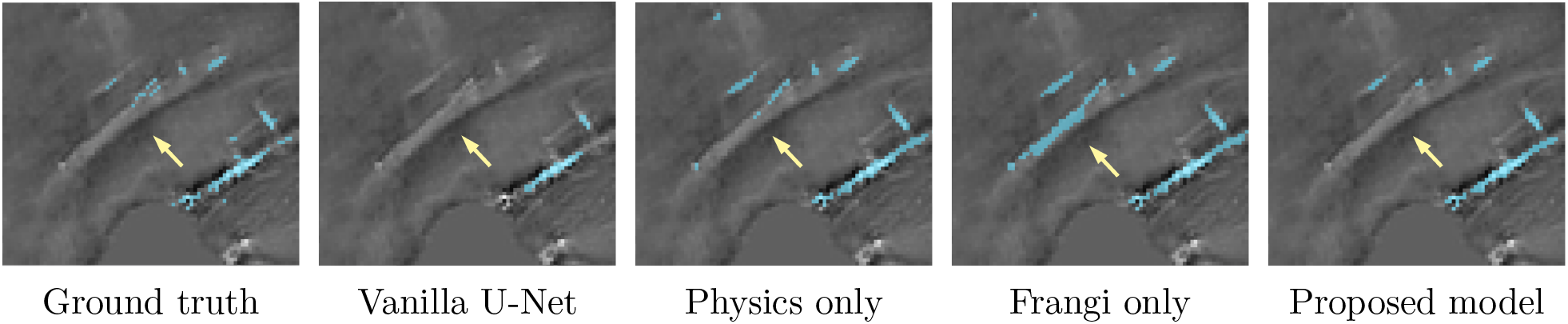
Qualitative ablation example illustrating the effect of the proposed physics and Frangi constraints on cerebral vein segmentation. From left to right: manual ground truth, vanilla U-Net, physics-only model, Frangi-only model, and the full proposed model. The Frangi-only model demonstrates increased sensitivity to tubular structures and improved recovery of small veins, but also introduces additional false-positive predictions. The full model provides the best balance between sensitivity and specificity, improving recovery and continuity of venous structures while limiting oversegmentation relative to the Frangi-only model.

An additional strength of the proposed framework is its robustness across multiple QSM reconstruction methods. Different QSM regularization strategies produce substantially different susceptibility contrast, noise texture, dynamic range, and artifact characteristics, all of which influence venous conspicuity and segmentation difficulty. Although Dice scores remained relatively stable across reconstruction methods compared to TGV-QSM, ASSD errors were similarly preserved, with MEDI and STAR-QSM achieving Dice scores comparable to TGV-QSM and similar or lower ASSD. L1 reconstruction also maintained relatively strong performance, whereas iLSQR showed the largest reduction in Dice and precision. Recall remained highly consistent across the alternative reconstruction methods, suggesting that reconstruction-specific differences primarily affected the specificity and spatial accuracy of the predicted venous maps rather than overall sensitivity. Consequently, training across multiple QSM reconstructions was important to ensure generalizability, improve robustness to reconstruction-specific appearance shifts, and reduce overfitting to a single susceptibility reconstruction style. In the present study, five QSM reconstruction methods were incorporated during training, exposing the network to substantial variability in venous contrast and artifact behaviour. Additional reconstruction approaches were explored but were ultimately excluded because severe degradation in venous contrast would have required extensive manual resegmentation. In contrast, the selected reconstruction methods preserved sufficient venous visibility such that the original TGV-QSM segmentations could be transferred with only minimal refinement, allowing efficient generation of multi-reconstruction training labels while still capturing meaningful variability in susceptibility appearance. Despite this improved robustness across reconstruction methods, segmentation performance still varied regionally, likely reflecting underlying differences in venous anatomy, vessel caliber, and local susceptibility contrast throughout the brain.

Regional analysis showed that segmentation performance was highest in the occipital lobe and lowest in the frontal lobe, reflecting differences in venous anatomy and regional susceptibility effects. Occipital regions contain larger, more continuous veins that are more readily resolved on QSM, whereas the frontal lobe is located in proximity to air-filled paranasal sinuses, which generate strong susceptibility gradients and may reduce venous contrast through residual background fields and QSM reconstruction artifacts [54, 55]. Diameter-stratified analysis further demonstrated that the primary benefit of the proposed framework lies in improved recovery of smaller cortical veins. Improvements in global overlap metrics were relatively modest because Dice similarity is dominated by larger venous structures that are already segmented reliably across deep learning approaches. In contrast, the largest gains were observed in vessels below 1,mm and 2,mm in diameter, indicating that the proposed constraints preferentially improve sensitivity to fine venous anatomy. This is further supported by improvements in clDice, which reflects vascular continuity beyond voxel-wise overlap alone. Performance was similar across ablations for veins larger than 4,mm, where segmentation is less challenging. A similar pattern was observed in the downstream OEF analyses, where the strongest correlations with GCM-derived OEF occurred in the parietal and occipital lobes, while weaker correlations were observed in the frontal and temporal lobes. The greater sensitivity of smaller cortical veins to partial-volume contamination and reduced SNR, together with regional susceptibility effects, may contribute to the lower reliability of venous susceptibility measurements in these regions.

The significant positive correlations observed between venous QSM-derived OEF and GCM-derived OEF are consistent with prior work supporting the physiological validity of susceptibility-based venous oximetry. Fan et al. [19] reported moderate agreement between venous QSM-OEF and calibrated fMRI-derived OEF in 7T imaging, while Kudo et al. [30] demonstrated significant correspondence between QSM-derived OEF and ^15^O-PET. However, these prior approaches primarily relied on larger cerebral veins and venous sinuses, where susceptibility measurements are less affected by partial-volume contamination and segmentation inaccuracies. In contrast, smaller cortical draining veins are expected to provide more spatially localized information regarding regional oxygen metabolism. An additional source of disagreement between venous QSM-derived OEF and GCM-derived OEF is that venous QSM relies on assumptions regarding hematocrit and the susceptibility contribution of deoxyhemoglobin, whereas GCM-derived OEF is inferred from calibrated BOLD and CBF responses to respiratory manipulation. Consequently, differences in the physiological sensitivity of the two models, particularly with respect to hematocrit and blood composition, may contribute to imperfect agreement between the techniques. Further, the pCASL acquisition used for GCM-derived OEF has lower spatial resolution than the GRE acquisition used for venous QSM, increasing partial-volume effects.

The present findings suggest that improved recovery of smaller venous structures may enhance the physiological sensitivity of venous QSM-OEF measurements. Notably, Fan et al. utilized high-resolution 7T imaging with substantially higher venous contrast and SNR than typically achievable at 3T [19]. In the current study’s 3T cohort, no significant correlation with GCM-derived OEF was observed using the MSVF approach, despite significant associations emerging with the proposed segmentation framework. These findings highlight the importance of more complete venous recovery, particularly within smaller cortical veins that are more susceptible to partial-volume effects and are often incompletely captured by conventional vessel filtering approaches. However, inclusion of smaller veins also increases susceptibility to noise, partial-volume effects, and reconstruction artifacts, particularly in lower-resolution 3T acquisitions with anisotropic voxel dimensions, which further reduce the visibility and boundary definition of small cortical veins. As a result, although sensitivity to physiological variation is improved, the increased variability observed within groups may partially reflect the inclusion of noisier veins. The difference in OEF between approaches is illustrated in Figure 6.

**Figure 6:**
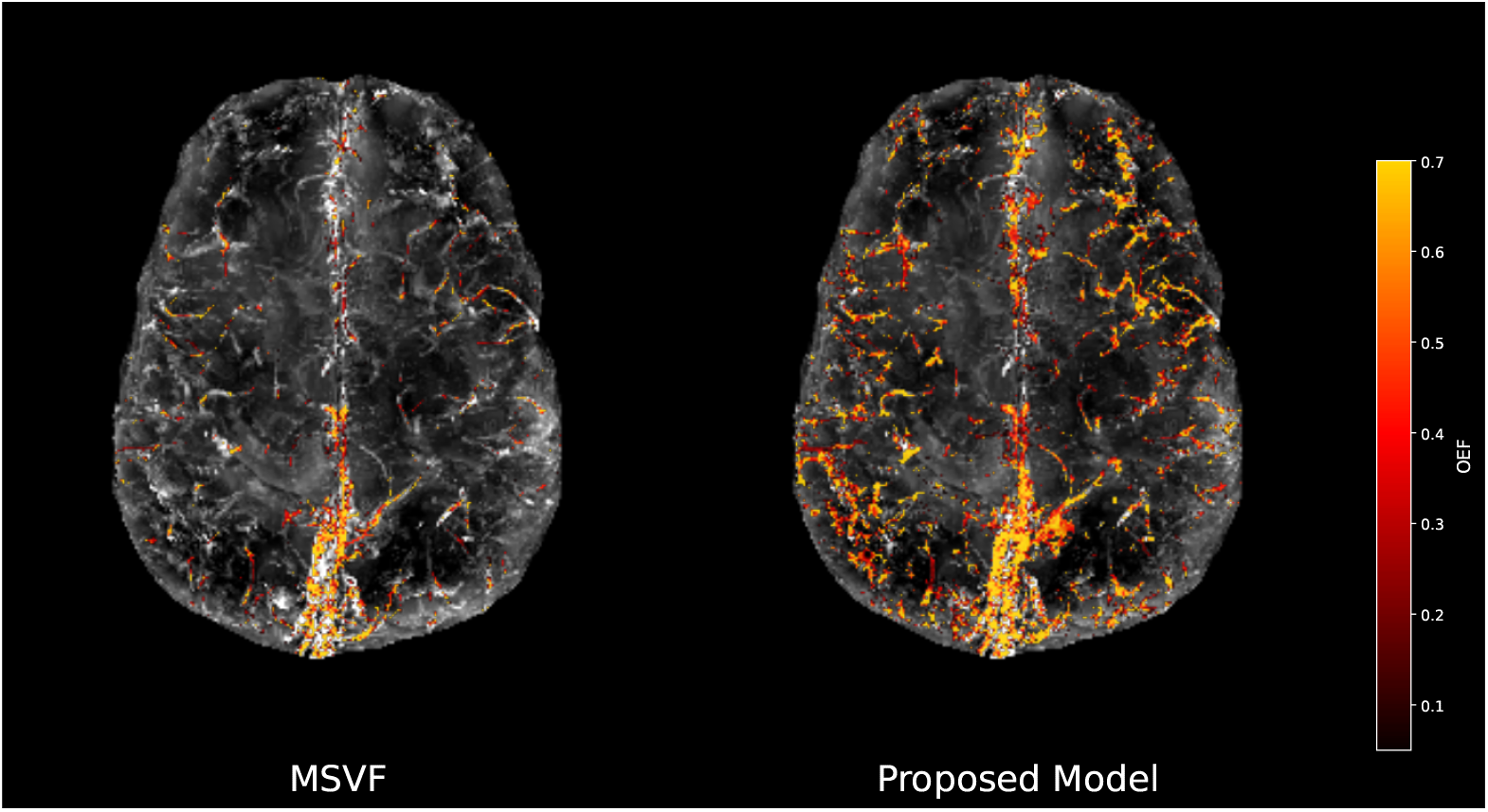
Maximum intensity projection comparison of venous OEF maps generated using the MSVF approach (left) and the proposed model (right). The proposed model demonstrates improved segmentation of smaller veins, leading to more spatially distributed venous OEF measurements compared to the more limited large-vessel emphasis of MSVF.

### 4.1 Limitations

Several limitations should also be considered when interpreting the present findings. Although the proposed constraints are intended to reduce dependence on manual annotations, the framework remains fundamentally data-driven and therefore still depends on the quality and consistency of the training labels. This is particularly relevant for cerebral vein segmentation, where fine cortical vessels are inherently difficult to delineate and some degree of labeling uncertainty is unavoidable. In addition, despite incorporating data from multiple centers and scanners, both 3T and 7T acquisitions, and subjects with conditions such as diabetes, CAD, and AD, the overall dataset remains relatively limited in size and variability. Larger and more heterogeneous datasets spanning broader scanner types and vendors, such as General Electric (GE) scanners, along with a wider range of acquisition protocols and pathological conditions, would likely improve both robustness and generalizability. Artifacts also remain a challenge, particularly in lower-resolution QSM acquisitions. Ringing and streaking artifacts can generate elongated susceptibility patterns that resemble tubular venous anatomy and may therefore be incorrectly classified as veins, especially in 3T data where venous contrast is reduced. Furthermore, OEF estimation in the present study relied on assumed physiological constants, including hematocrit, rather than subject-specific measurements. Inter-subject variability in hematocrit and blood composition would therefore introduce bias into the absolute OEF estimates, particularly in aged populations with vascular or metabolic disease.

Beyond venous segmentation, the proposed framework may also have broader applications in quantitative MRI. The physics-constrained approach encourages predictions that are consistent with the expected field perturbations and morphology of venous susceptibility sources. Improved characterization of venous susceptibility may facilitate more accurate estimation of non-vascular susceptibility components in techniques such as *χ*-separation, where venous signal can act as a confounding paramagnetic source [56]. Due to veins representing a major paramagnetic contributor to QSM contrast, improved delineation of venous anatomy may help disentangle vascular susceptibility effects from other sources such as iron deposition and myelin [16]. Generally, incorporating anatomy-aware and physics-informed priors into data-driven frameworks may provide a useful strategy for improving the robustness and optimization of QSM reconstruction methods across varying field strengths and pathological conditions.

## 5 Conclusions

The proposed framework improves QSM-based cerebral vein segmentation by combining data-driven learning with susceptibility physics and vessel morphology priors. The results suggest that these constraints are particularly useful for improving sensitivity to smaller cortical veins, which may in turn enhance the physiological sensitivity of venous QSM-derived OEF measurements. These findings support the potential of susceptibility-based venous imaging as a scalable and non-invasive approach for studying cerebral oxygen metabolism from routinely acquired GRE/QSM data.

## Acknowledgments

The authors thank all participants for their time and cooperation. The authors also thank the staff at the Montreal Heart Institute, the Max Planck Institute, the Montreal Neurological Institute, the Douglas Hospital, and the Centre de recherche de l’Institut universitaire de gériatrie de Montréal for their assistance with data acquisition. The authors also thank Habiba Megawer, Sophie Courtemanche, and Tanusha Kanagasingam for additional help with manually segmenting veins.

## Author Contributions

YCA. and AR contributed equally to this work. YCA and AR designed and implemented the deep learning framework, and wrote the manuscript. VR contributed to data processing, evaluation, and writing. NS, HG, contributed to manual vein segmentations. NS and HG for providing manual segmentations of veins. YX contributed to project conceptualization, data validation, and writing. BPT contributed to validation and writing. CJG supervised the project, contributed to study design, funding, data acquisition and writing of the manuscript.

## Supporting Information

### A Training Hyperparameters

#### A.1 Optimisation

**Table S1:** Training and optimisation hyperparameters.

| Hyperparameter | Value |
| --- | --- |
| Optimiser | SGD (Nesterov) |
| Initial learning rate | $10^{-2}$ |
| Momentum | 0.99 |
| Weight decay | $3 \times 10^{-5}$ |
| Learning rate schedule | Polynomial decay to 0 |
| Epochs | 500 |
| Batch size | 2 |
| Patch size | $128 \times 160 \times 112$ voxels |
| Voxel spacing | 0.6 mm isotropic |
| Foreground oversample | 33% |
| Deep supervision | Enabled ($w_i = 1/2^i$) |

#### A.2 Loss Function

The total loss was defined as

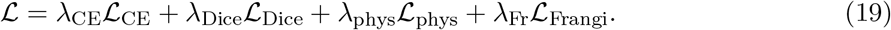

**Table S2:** Top-level loss weights.

| Loss term | Weight |
| --- | --- |
| Cross-entropy ($\mathcal{L}_{\text{CE}}$) | 1.0 |
| Soft Dice ($\mathcal{L}_{\text{Dice}}$) | 1.0 |
| Physics loss ($\mathcal{L}_{\text{phys}}$) | 2.0 |
| Frangi loss ($\mathcal{L}_{\text{Frangi}}$) | 0.25 |

##### A.2.1 Physics field-matching loss

The physics loss consisted of four sub-terms:

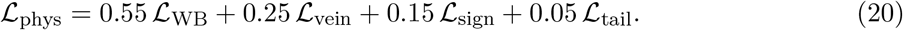

The venous susceptibility value *χ*_blood_ was estimated separately for each field strength and reconstruction method from the training data. The tail term used the top 10% of field residuals.

**Table S3:** Full model segmentation performance across cortical lobes, averaged across *n* = 10 held-out cases (TGV-QSM, mean ± std). The Frangi-only model demonstrates increased sensitivity to tubular structures and improved recovery of small veins, but also introduces additional false-positive predictions. The full model provides the best balance between sensitivity and specificity, improving recovery and continuity of venous structures while limiting oversegmentation relative to the Frangi-only model.

| Region | DSC | Precision | Recall | ASSD (mm) |
| --- | --- | --- | --- | --- |
| Frontal Lobe | $0.58 \pm 0.06$ | $0.54 \pm 0.09$ | $0.63 \pm 0.04$ | $1.62 \pm 0.75$ |
| Temporal Lobe | $0.68 \pm 0.06$ | $0.61 \pm 0.09$ | $0.78 \pm 0.03$ | $0.83 \pm 0.33$ |
| Parietal Lobe | $0.73 \pm 0.06$ | $0.76 \pm 0.04$ | $0.70 \pm 0.08$ | $0.69 \pm 0.27$ |
| Occipital Lobe | $0.79 \pm 0.06$ | $0.71 \pm 0.08$ | $0.90 \pm 0.01$ | $0.38 \pm 0.20$ |

##### A.2.2 Frangi vesselness loss

For the image branch, vesselness was computed at scales

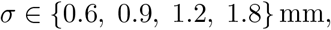

with Frangi parameters

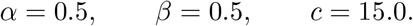

For the prediction branch, scales

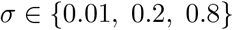

were used with

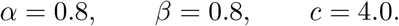

The vesselness gate used a sigmoid function with sharpness 6.0 and midpoint *τ* = 0.15.

### B Regional Performance

Regional performance of the full model across cortical lobes is summarized in Table S3. The highest performance is observed in the occipital lobe, which also exhibits the lowest ASSD. The parietal and frontal lobes show comparable performance, while the temporal lobe yields lower overlap and higher surface distance error.

Regional correlations between venous QSM-derived OEF and GCM-derived OEF are shown in Supplementary Figures S1–S2. The proposed framework demonstrated the strongest associations in the parietal and occipital lobes, with a significant whole-brain correlation also observed. In contrast, OEF derived using the MSVF approach showed weaker and non-significant associations across all regions.

**Figure S1:**
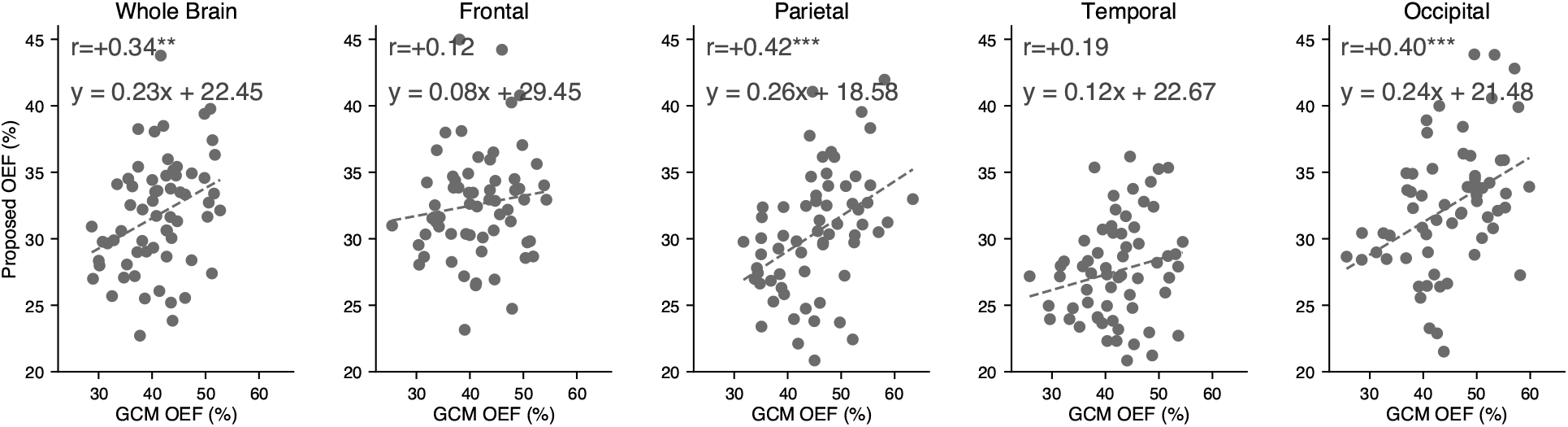
Regional correlations between venous OEF derived using the proposed segmentation framework and OEF estimated using the gas calibration model (GCM). Scatter plots are shown for whole-brain, frontal, parietal, temporal, and occipital regions. The strongest associations were observed in the parietal and occipital lobes, while weaker correlations were observed in the frontal and temporal lobes.

**Figure S2:**
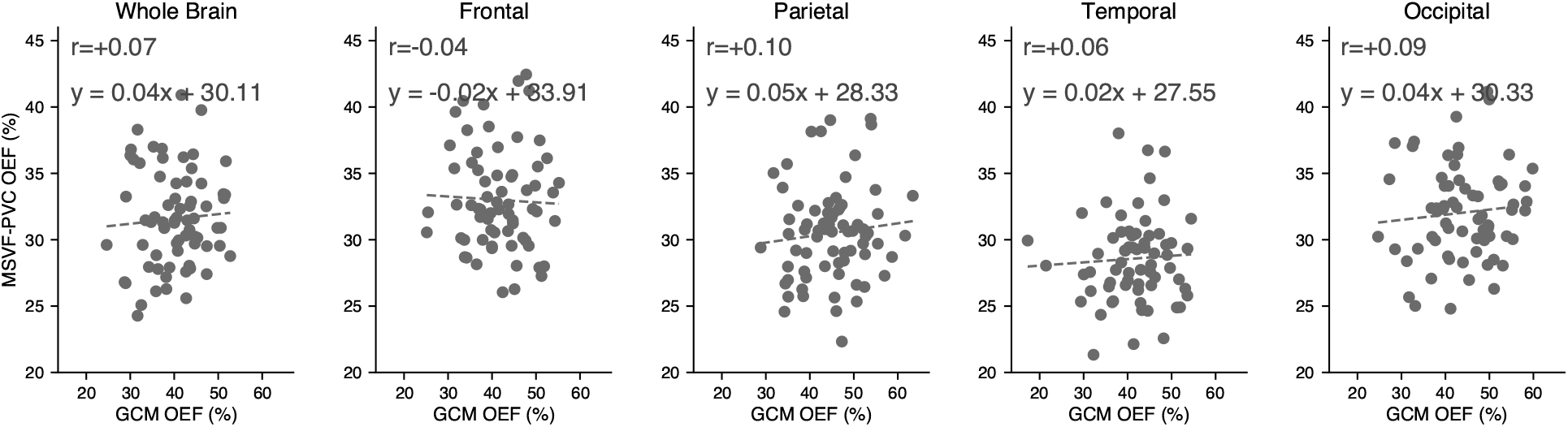
Regional correlations between venous OEF derived using the multiscale vessel filtering (MSVF) approach and OEF estimated using the gas calibration model (GCM). Scatter plots are shown for whole-brain, frontal, parietal, temporal, and occipital regions. Compared with the proposed framework, MSVF-derived OEF demonstrated weaker and largely non-significant associations across all regions.

